# The C-terminal regulatory domain of IPMS enzyme maintains leucine homeostasis by bypassing a hidden negative feedback loop in plants

**DOI:** 10.1101/2024.06.14.598950

**Authors:** Mohan Varghese, Roshan Kumar, Aprajita Sharma, Asif Lone, Jonathan Gershenzon, Naveen C Bisht

## Abstract

In the leucine biosynthesis pathway, homeostasis is achieved through a feedback regulatory mechanism facilitated by binding of the end-product Leu at the C-terminal regulatory domain of the first committed enzyme, isopropylmalate synthase (IPMS). *In*-*vitro* studies showed that removal of the regulatory domain abolishes the feedback regulation on plant IPMS while retaining its catalytic activity. However, the physiological consequences and underlying molecular regulation upon removal of the IPMS C-terminal domain on the Leu flux have not been previously explored in plants. Here, we show that in the absence of its regulatory domain, an unexpected alternative regulatory loop acts to control plant IPMS catalysis. Removal of IPMS regulatory domain using CRISPR/Cas9 significantly reduced the formation of end-product Leu *in*-*planta*, but increased the levels of Leu pathway intermediates. Additionally, delayed growth was observed when IPMS devoid of regulatory domain was introduced into *IPMS*-null mutants of E. *coli* and *Arabidopsis*. Combining the metabolomic and biochemical analysis, we found that the Leu pathway intermediate, α-ketoisocaproate, was a competitive inhibitor of IPMS with a truncated regulatory domain. Thus, we demonstrate that the C-terminal regulatory domain of IPMS is biologically favored since it maintains Leu homeostasis while bypassing the possibility of competitive inhibition by a pathway intermediate.

**Significance:** Till date it was known that the limited profile of essential amino acid-leucine in plants, is majorly due to the feedback inhibition of its pathway enzyme, IPMS, by the binding of accumulating leucine at its regulatory domain. So, can we increase the leucine pool in plants by removing the IPMS regulatory domain? Herein we show that, the targeted removal of this domain under native condition had led to low leucine pool and compromised growth phenotype but observed an accumulation in the leucine pathway intermediates. We uncover a hidden function of the IPMS regulatory domain to avoid an intermediate inhibition on IPMS activity, which could limit the end-product. The study brings an unknown regulatory checkpoint in maintaining leucine homeostasis in plants.

## Introduction

The existence of allosteric enzymes in metabolic pathways is meant to regulate the accumulation of an end-product by acting as a rate-limiting enzyme. Typically, the accumulating end-product binds to a regulatory domain found in these allosteric enzymes to regulate their enzyme activity (Galili et al., 2016; Koon et al., 2004). Such a feedback inhibition loop is typical in the anabolic pathways of amino acids which at the cellular level enables plants to ensure that the nitrogen and carbon resources for their synthesis are used only when supplies are low. Concurrently, this feedback loop is one of the major reasons for the limited content of amino acids like leucine (Leu) in crop plants, which are essential for humans and animals (Galili et al., 2016). Leu plays a very important role in overall human health, including *de novo* biosynthesis of muscle fibers, muscle repair, preventing muscle loss during aging, regulating blood sugar levels, and stimulating brain functions and immunity (Monirujjaman and Ferdouse, 2014). In the Leu biosynthesis pathway of plants, the activity of its first committed enzyme, isopropylmalate synthase (IPMS; EC 2.3.3.13), is known to be tightly regulated allosterically by the end-product Leu through its C-terminal (C-ter) regulatory domain.

IPMS exists at the branch point between Leu and valine (Val) biosynthesis and so regulates flux in both pathways. It redirects a common Leu-Val substrate, 3-methyl-2-oxobutanoate (a.k.a 2-oxoisovalerate, 2-OIV) towards Leu biosynthesis by catalyzing an aldol-type condensation between 2-OIV and acetyl-CoA, yielding 2-isopropylmalate (2-IPM) (**Figure 1A**). 2-IPM is then converted to 3-isopropylmalate (3-IPM) by isopropylmalate isomerase (IPMI; EC 4.2.1.33). The ultimate intermediate, one-carbon elongated 4-methyl-2-oxopentanoate (a.k.a α-ketoisocaproate, KIC) is formed by the oxidative decarboxylation of 3-IPM by isopropylmalate dehydrogenase (IPMDH; EC 1.1.1.85). KIC then undergoes a reversible transamination catalyzed by a branched-chain aminotransferase (BCAT; EC 2.6.1.42) to yield the end-product Leu in the chloroplast. When produced in sufficient quantity, Leu binds at the IPMS C-ter regulatory domain limiting the intake of 2-OIV into the IPMS active site and thus directing the flux towards Val (Field et al., 2006; de Kraker et al., 2007). Thus, the C-ter regulatory domain of IPMS facilitates the regulation of both Leu and Val biosynthesis directly and indirectly, respectively. Hence, targeting the IPMS regulatory domain provides a framework for investigating the influence of this enzyme on cellular metabolism.

**Figure 1:**
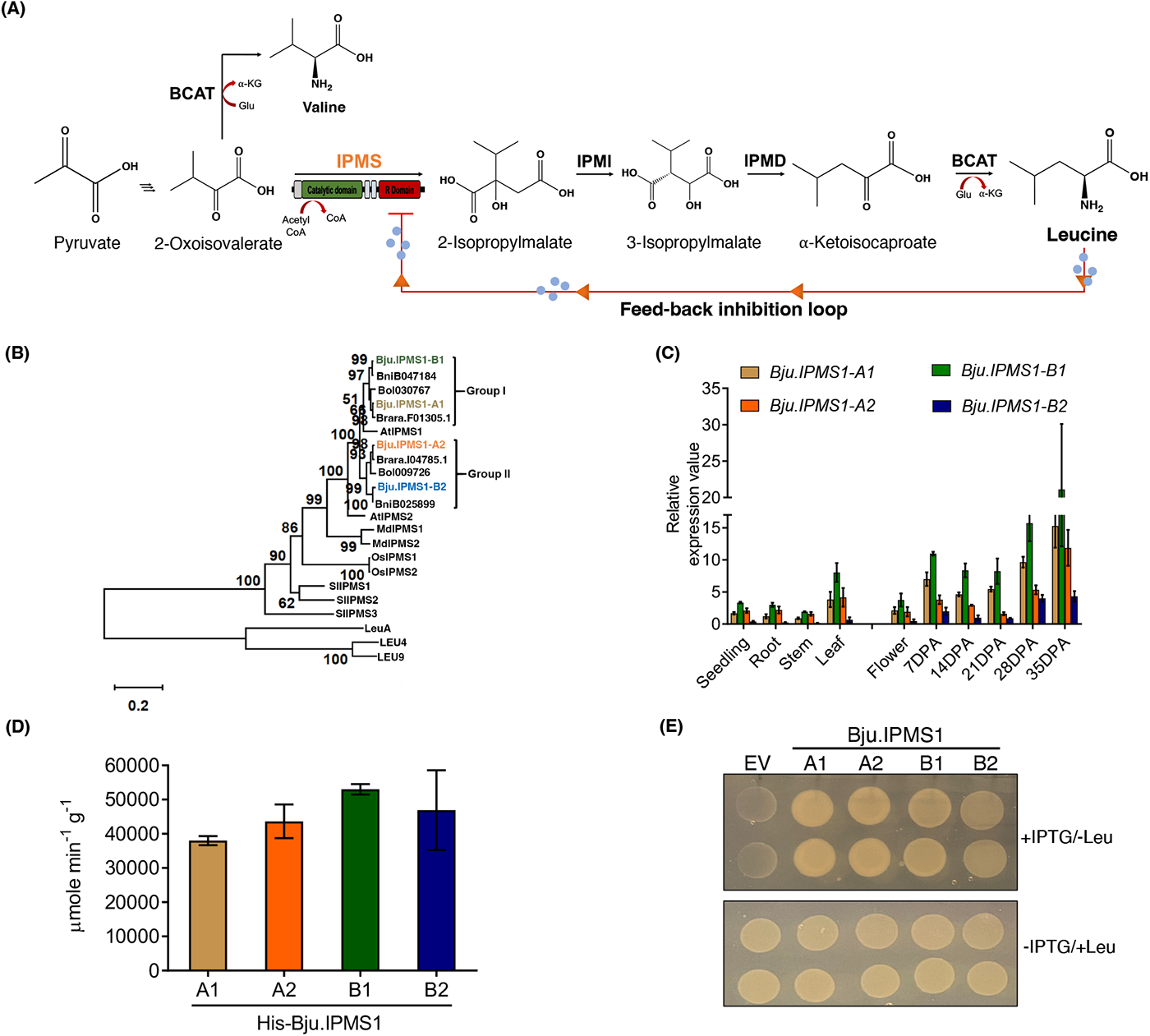
Molecular characterization of four *IPMS1* homologs from *Brassica juncea*. **(A)** Leu biosynthesis pathway. IPMS acts as a branch-point enzyme with the Val pathway that also regulates Leu biosynthesis by feedback inhibition. **(B)** Phylogenetic analysis of *B. juncea* IPMS1 (A1, A2, B1, B2) proteins and those of other plant species. The branch number represents a percentage of the bootstrap values in 1000 sampling replicates, and the scale bar indicates branch lengths. The IPMS of *Mycobacterium* (LeuA) and yeast (LEU4 and LEU9) were used as out-group proteins. **(C)** Transcript profiling of the *Bju.IPMS1* homologs in vegetative and reproductive tissue types. Real-time PCR amplifications were performed for each target gene in two biological replicates with two technical replicates each. Error bars represent the standard error of the mean. The expression of *BjuActin* was used to normalize the data (set at 100). **(D)** Spectrophotometric end-point assay of four recombinant Bju.IPMS1 proteins with 5,5′-dithiobis-(2-nitrobenzoic acid) to show *in vitro* activity. Data represent mean ±SD (n=8). **(E)** Demonstration of the activity of the *B. juncea* IPMS proteins by the rescue of an *IPMS*-null *E. coli* strain. *E. coli* CV512 (DE3) harboring the four *pET28a::Bju.IPMS1* CDS separately or an empty vector (pET28a) control was grown on M9 minimal media with 1 mM IPTG (+IPTG/-Leu) or M9 minimal media with 2 mM Leucine (-IPTG/+Leu) and growth was photographed after 72 hours. Assays were performed in three biological replicates, with similar results.

While studies on plant IPMSs to date have proved their involvement in Leu biosynthesis as well as in other plant traits and signaling, our understanding of the biochemical regulation of this enzyme is quite sparse (de Kraker et al., 2007; He et al., 2019; Cao et al., 2019). In the model plant *Arabidopsis thaliana*, a random point mutation in the C-ter regulatory domain provides a partially relaxed Leu feedback-resistant mutant of IPMS (Xing & Last, 2017). Studies in the bacteria *Neisseria meningitidis* (NmeIPMS) and *Mycobacterium tuberculosis* (MtIPMS) show that the C-ter regulatory domain of IPMS also contributes to its catalytic ability since the removal of the domain dramatically reduces IPMS activity (Huisman et al., 2012). On the contrary, *in vitro* study of the *Arabidopsis* IPMS shows that the removal of the C-ter regulatory domain abolished Leu inhibition, but did not otherwise affect activity (deKraker and Gershenzon, 2011). Nonetheless, the impact of the plant IPMS C-ter regulatory domain at the metabolic level is unexplored *in vivo*.

Further reports on plant IPMS points to the evolutionary recruitment of the enzyme into specialized metabolism, like MAMS in glucosinolate biosynthesis in Brassicaceae (deKraker and Gershenzon, 2011), SlIPMS3 in acylsugar metabolism in trichomes of tomatoes (Ning et al., 2015), MdCMS in citramalate biosynthesis for ester formation in apples (Sugimoto et al., 2021) and PcIBMS1 in pogostone biosynthesis in *Pogostemon cablin* (Wang et al., 2022). One of the major modifications that facilitated the neo-functionalism of these IPMS-like enzymes is the deletion of the regulatory domain from the C-ter region of IPMS rather than specific modification of Leu binding residues in the C-ter domain, showing its specific role in Leu biosynthesis (deKraker and Gershenzon, 2011).

These prior studies on the significance of the C-ter regulatory domain for end-product regulation of IPMS and its importance in Leu biosynthesis raise a few important questions. First, (1) what is the physiological consequence of removing the IPMS C-ter regulatory domain on Leu-Val metabolic flux, and (2) what is the significance of this domain in maintaining Leu homeostasis in plants? The answers will add to the limited information on IPMS regulation and could facilitate engineering of the amino acid profile in crop plants. Therefore, we developed IPMS C-ter edited mutant lines in the globally cultivated oilseed crop *Brassica juncea* using CRISPR/Cas9 and also overexpressed the C-ter truncated IPMS in *IPMS*-null *Arabidopsis* and *E. coli*. Using a combination of genetic, metabolomic, biochemical and phenotypic analysis, we showed that, in addition to its regulatory role, the IPMS C-ter regulatory domain maintains Leu flux by avoiding possible competitive inhibition by the immediate precursor of Leu.

## Results

### *B. juncea* genome encodes four functional IPMS proteins

Previously, in *A. thaliana* two *IPMS* genes were characterized both *in vitro* and *in vivo* for their role in Leu biosynthesis (deKraker 2007). To identify the corresponding *IPMS* homologs in *B. juncea, AtIPMS1* (At1g18500) and *AtIPMS2* (At7g14040) were used as BLAST queries to the *B. juncea* genome, then available at BRAD database (http://brassicadb.org/brad/). A total of four homologs, viz, *BjuA022211, BjuA044305, BjuB029278* and *BjuB032394* at chromosomes A06, A09, B04 and B03, respectively were identified in the allotetraploid *B. juncea* and at least two sequences in other diploid *Brassica* species **(Supplementary Table S1)**.

Phylogenetic analysis using the identified IPMS protein sequences from Brassicaceae and other plant species clustered all four *B. juncea* IPMS encoding genes (*Bju.IPMS*) with the *AtIPMS1* and grouped them into two distinct subclades viz., the orthologous group I and group II, suggesting duplication of *IPMS* genes in the *Brassica* lineage (**Figure 1B**). Based on close phylogeny with AtIPMS1, the four *B. juncea IPMS* homologs were designated as *Bju.IPMS1-A1 (BjuA022211)* and *Bju.IPMS1-B1 (BjuB029278)* from orthologous group I, and *Bju.IPMS1-A2 (BjuA044305)* and *Bju.IPMS1-B2 (BjuB032394)* from orthologous group II. The encoded Bju.IPMS1 protein sequences share 84% to 91% identity with the *Arabidopsis* counterpart and 90% to 96% identity between each other. The predicted amino acid sequences of all four Bju.IPMS1 harbor the N-terminal conserved (*⍺/β*)_8_ barrel structure as a catalytic domain and a ∼150 aa regulatory domain at its C-ter end, as also reported for the AtIPMS1 and LeuA **(Supplementary Figure S1).**

The multiplicity of *IPMS1* homologs led us to determine their expression pattern across different developmental stages of *B. juncea*, using qRT–PCR analysis. All four *Bju.IPMS1* homologs were ubiquitously expressed, with a higher accumulation of their transcripts during the reproductive stage compared to the vegetative phase (**Figure 1C**).

Further, to explore their *in vitro* enzyme characteristics, the enzyme activity of the purified recombinant Bju.IPMS1 proteins was assessed using a spectrophotometric end-point assay with 5,5-dithio-bis-(2-nitrobenzoic acid) (DTNB) (**Supplementary Figure S2, Figure 1D**). The assay showed that all four Bju.IPMS1 enzymes can catalyze the aldol condensation between acetyl CoA and 2-oxoisovalerate (2-OIV). To get detailed kinetic parameters, assays were carried out with varied concentrations of substrates (2-OIV, 50 μM to 4000 μM; Acetyl CoA, 10 μM to 2000 μM) (**Table 1**). Kinetic analysis using 2-OIV and acetyl CoA revealed that the four Bju.IPMS1 proteins have comparable catalytic efficiency (*k*_cat_/*K*_m_) for 2-OIV, suggesting that all four Bju.IPMS1 enzymes contribute to Leu biosynthesis in an indistinguishable manner.

**Table 1.**
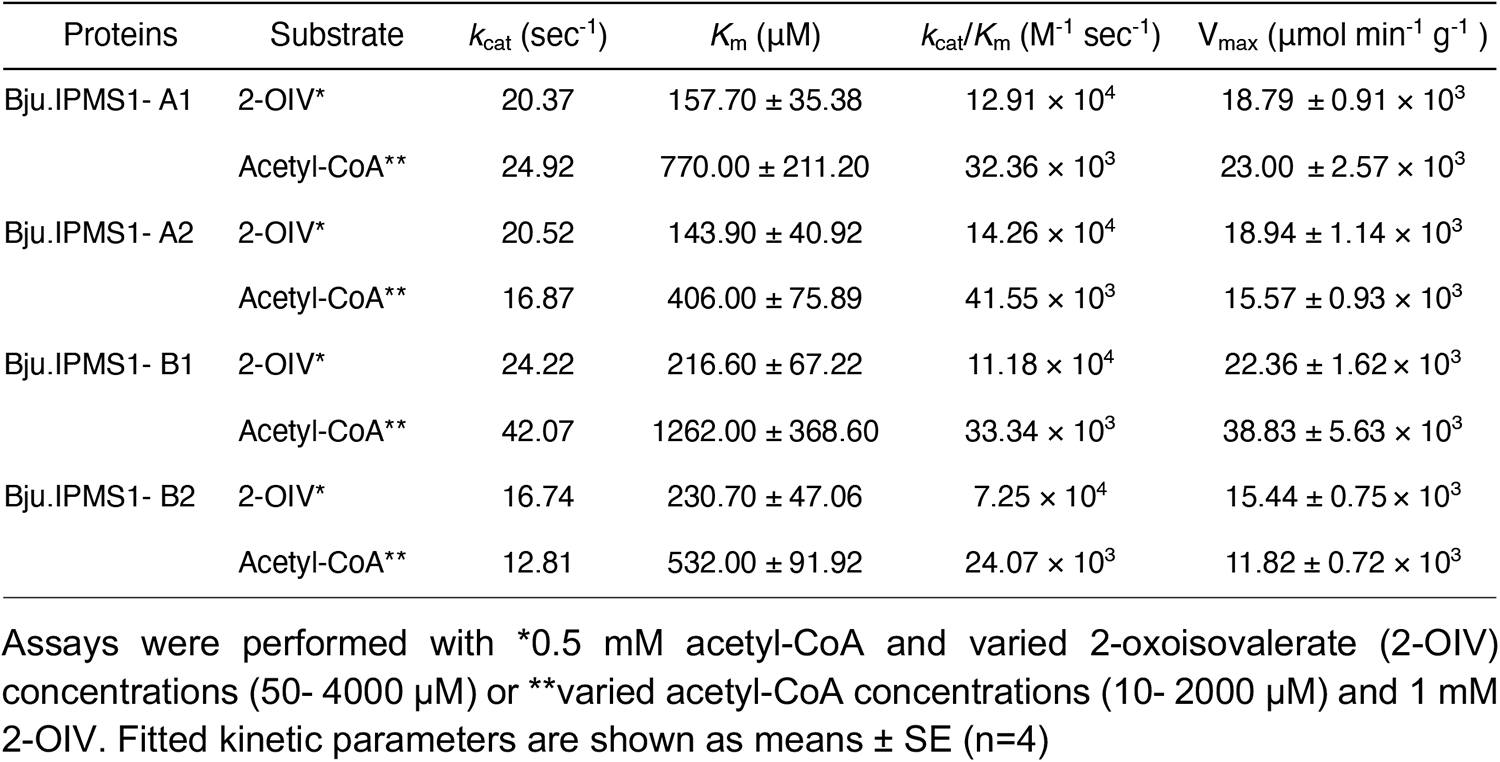
Kinetic parameters of the recombinant Bju.IPMS1 proteins.

| Proteins | Substrate | $k_{\text{cat}}$ (sec<sup>-1</sup>) | $K_m$ (μM) | $k_{\text{cat}}/K_m$ (M<sup>-1</sup> sec<sup>-1</sup>) | $V_{\text{max}}$ (μmol min<sup>-1</sup> g<sup>-1</sup>) |
| --- | --- | --- | --- | --- | --- |
| Bju.IPMS1- A1 | 2-OIV* | 20.37 | 157.70 ± 35.38 | 12.91 × 10<sup>4</sup> | 18.79 ± 0.91 × 10<sup>3</sup> |
|  | Acetyl-CoA** | 24.92 | 770.00 ± 211.20 | 32.36 × 10<sup>3</sup> | 23.00 ± 2.57 × 10<sup>3</sup> |
| Bju.IPMS1- A2 | 2-OIV* | 20.52 | 143.90 ± 40.92 | 14.26 × 10<sup>4</sup> | 18.94 ± 1.14 × 10<sup>3</sup> |
|  | Acetyl-CoA** | 16.87 | 406.00 ± 75.89 | 41.55 × 10<sup>3</sup> | 15.57 ± 0.93 × 10<sup>3</sup> |
| Bju.IPMS1- B1 | 2-OIV* | 24.22 | 216.60 ± 67.22 | 11.18 × 10<sup>4</sup> | 22.36 ± 1.62 × 10<sup>3</sup> |
|  | Acetyl-CoA** | 42.07 | 1262.00 ± 368.60 | 33.34 × 10<sup>3</sup> | 38.83 ± 5.63 × 10<sup>3</sup> |
| Bju.IPMS1- B2 | 2-OIV* | 16.74 | 230.70 ± 47.06 | 7.25 × 10<sup>4</sup> | 15.44 ± 0.75 × 10<sup>3</sup> |
|  | Acetyl-CoA** | 12.81 | 532.00 ± 91.92 | 24.07 × 10<sup>3</sup> | 11.82 ± 0.72 × 10<sup>3</sup> |

Assays were performed with \*0.5 mM acetyl-CoA and varied 2-oxoisovalerate (2-OIV) concentrations (50- 4000 μM) or \*\*varied acetyl-CoA concentrations (10- 2000 μM) and 1 mM 2-OIV. Fitted kinetic parameters are shown as means ± SE (n=4)

Additionally, to validate the IPMS function of *Bju.IPMS1 in vivo*, the *pET28a::Bju.IPMS1* gene constructs were transformed individually into an *IPMS*-null *E. coli* strain CV512 (DE3), and isopropyl-*β*-galactoside (IPTG) was used to induce the transcription of these constructs (**Figure 1E**). All four *Bju.IPMS1* constructs were able to complement the CV512 (DE3). Overall, the combined molecular and biochemical analysis suggests that the four *Bju.IPMS1* homologs share a close evolutionary relationship with plant *IPMS1*, while maintaining a high degree of transcript expression and IPMS activity.

### The Bju.IPMS1 enzyme devoid of its regulatory domain retains IPMS catalytic activity while exhibiting no feedback inhibition by Leu *in vitro*

Prior to *in vivo* editing of the C-ter regulatory domain of *B. juncea* IPMS1s, we checked the feedback inhibition of a candidate Bju.IPMS1-A1 by the end-product Leu *in vitro* and whether its IPMS activity is retained after the removal of C-ter regulatory domain. The recombinant Bju.IPMS1-A1 protein C-ter truncated until the end of sub-domain II (Asp 474), designated as Bju.IPMS1-A1ΔR, was purified over an Ni-NTA agarose affinity column and the DTNB enzyme assay was performed. The *in vitro* assay showed that the IPMS activity is retained in the Bju.IPMS1-A1ΔR variant with slightly higher catalytic activity than the full-length Bju.IPMS1-A1 (**Figure 2A**). Further, the DTNB assay performed in the presence of Leu at a series of concentrations from 0.25 mM to 5 mM showed that Bju.IPMS1-A1ΔR was insensitive to Leu concentrations up to 5 mM, whereas the activity of the full-length Bju.IPMS1-A1 enzyme was reduced to 60.5% at 0.25 mM and to 41.8% at the highest tested Leu concentration of 5 mM (**Figure 2B**). These *in vitro* data clearly suggest that the removal of the C-ter regulatory domain makes the enzyme insensitive towards allosteric feedback inhibition by the end-product Leu, but does not compromise Bju.IPMS1 catalysis towards 2-OIV.

**Figure 2.**
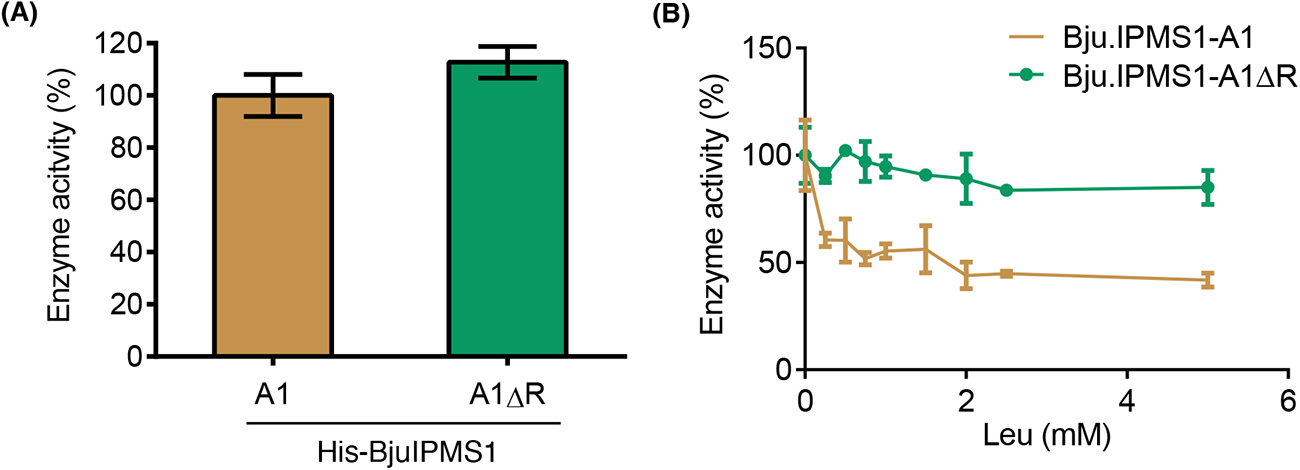
Effect of removing C-terminal regulatory domain on Bju.IPMS1-A1 activity and its feedback inhibition by Leu. **(A)** The activity of the regulatory domain truncated variant (Bju.IPMS1-A1ΔR) as a percentage of the full-length Bju.IPMS1-A1 protein set to 100%. **(B)** Effect of Leu inhibition on the enzymatic activities of Bju.IPMS1-A1ΔR variant and full-length Bju.IPMS1-A1 towards 2-oxoisovalerate (2-OIV) substrate. Activity of both full-length and truncated proteins was set to 100 % in the absence of Leu. Data represent mean ± SD (n = 4).

### CRISPR/Cas9-based editing of the *Bju.IPMS1* regulatory domain in *B. juncea* leads to reduced Leu level and compromised growth

To investigate the consequence of the catalytically active Bju.IPMS1ΔR protein (**Figure 2**) on Leu metabolic flux *in planta*, we designed the gRNAs from the 10^th^ exon, the start of the C-ter regulatory domain and transformed the SpCas9-sgRNAs binary vectors in *B. juncea* to specifically truncate the C-ter regulatory domain containing regions of *Bju.IPMS1*-*A1* and -*A2* genes (**Figure 3A and 3B**). Segregation of Cas9-induced mutations in the T2 advanced generation generated two C-ter edited Bju.IPMS1ΔR homozygous mutants, viz, *a_1_a_1_*ΔR and *a_2_a_2_*ΔR (**Figure 3C and 3D**). Each *a_1_a_1_ΔR* and *a_2_a_2_ΔR* mutant harbored a biallelic +1 insertion in the C-ter region of *Bju.IPMS1-A1* and -*A2,* respectively (**Figure 3C and 3D; Supplementary Figure S3**). Sequence analysis of the edited lines showed that the C-ter regulatory domain comprising 148 amino acids was effectively truncated by the gRNA to form the C-ter edited Bju.IPMS1ΔR proteins (**Figure 3E; Supplementary Text 1**). While *a_1_a_1_ΔR* and *a_2_a_2_ΔR* biallelic single mutants each were viable, we were unable to recover double (*a_1_a_1_ΔR* + *a_2_a_2_ΔR*) and any higher order mutants in T2 generation (**Supplementary Table S2**).

**Figure 3.**
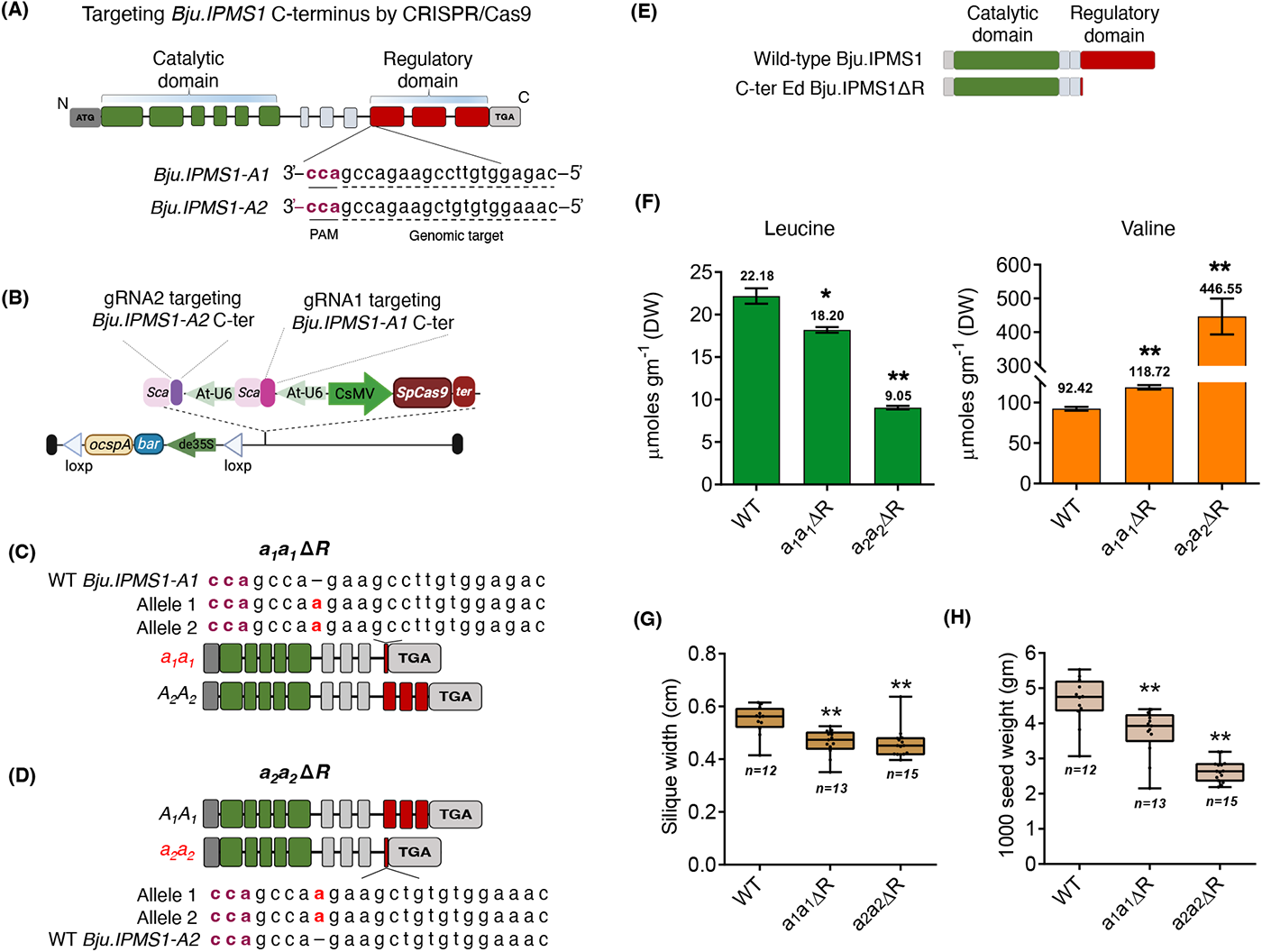
Effect of CRISPR/Cas9 editing of the Bju.IPMS1 C-terminal regulatory domain on Leu-Val flux *in vivo* and reproductive yield. (A) Schematic representation of *Bju.IPMS1* homologs (*A1* and *A2)* showing the PAM sites (maroon) and the genomic target sites (dotted lines) in the C-terminal regulatory domain for the respective gRNAs. **(B)** T-DNA map of *Bju.IPMS1* editing construct expressing *gRNA1* and *gRNA2* under individual At-U6 promoters and *SpCas9* under CsMV promoter. Selection of two Bju.IPMS1ΔR edited lines based on the mutations in the C-ter regulatory domains viz., (C) *a1a1*ΔR mutant harbouring homozygous *Bju.IPMS1-A1* C-ter mutated alleles (*a1a1*) and wild-type *Bju.IPMS1-A2*; and (D) *a2a2*ΔR mutant harbouring homozygous *Bju.IPMS1-A2* C-ter mutated alleles (*a2a2*) and wild-type *Bju.IPMS1-A1*. Indels in the CRISPR/Cas9-generated *Bju.IPMS1-A1* and *-A2* genes leading to premature stop codon are highlighted as red bold. **(E)** Schematic linear representation of the full-length Bju.IPMS1 wild-type protein and the C-ter Ed Bju.IPMS1ΔR, produced in the two mutants due to premature translation stop codon. **(F)** Free Leu and Val levels in one month old leaves of *a1a1*Δ*R* and *a2a2*Δ*R* mutants. The amino acid profile was analysed using LC-MS/MS QTRAP 6500+ in T3 generation plants and values (μmoles gm^-1^ dry weight) are expressed as mean ± SE (n=3). Data are summarized in Supplemental Table 3. Reproductive parameters of the two C-ter Ed Bju.IPMS1ΔR mutants from T3 generation show reduced **(G)** silique width and **(H)** 1000 seed weight. Asterisks in **F-H** indicate significant differences compared to the wild-type *B. juncea* variety Varuna (*, P < 0.05 and **, P < 0.01 Student’s t-test).

We then analyzed the Leu and Val concentrations in one-month old leaves of the C-ter edited Bju.IPMS1ΔR mutants using LC-MS/MS based quantification. While we anticipated that such mutants would have increased levels of end-product Leu; to our surprise both *a_1_a_1_ΔR* and *a_2_a_2_ΔR* mutants had 0.82-and 0.40-fold reduced Leu content, respectively (18.20 and 9.05 μmoles g^-1^ DW, respectively) compared to the wild-type plant having 22.19 μmoles g^-1^ DW Leu (**Figure 3F; Supplementary Table S3**). The mutants further have a concomitant 1.28-(*a_1_a_1_ΔR*) and 4.83-(*a_2_a_2_ΔR)* fold increase in Val content (118.73 and 446.55 μmoles g^-1^ DW respectively) in leaves compared to 92.42 μmoles g^-1^ DW Val in the wild-type plant (**Figure 3F**). The concurrent reduction and increase in Leu and Val profiles, respectively in the C-ter edited Bju.IPMS1ΔR lines of *B. juncea* revealed that the regulatory domain of IPMS has a great significance on Leu-Val metabolomic flux.

Furthermore, to investigate the consequence of the reduced Leu content in C-ter edited Bju.IPMS1ΔR mutants, we measured and analysed major agronomic parameters like plant height, silique length, silique width, and seed weight (**Figure 3G & H; Supplementary Figure S4; Supplementary Table S4**). The data showed that major reproductive traits like silique architecture and seed weight were significantly affected in the *a_1_a_1_ΔR* and *a_2_a_2_ΔR* mutants. Silique width and seed biomass in C-ter edited Bju.IPMS1ΔR mutants were significantly reduced by 17% and 45%, respectively, compared to the wild-type plant (**Supplementary Table 4**). Overall, the metabolic and phenotype results show that the C-ter regulatory domain of the Bju.IPMS1 enzyme has a key role in maintaining the Leu-Val flux and facilitated normal plant growth.

### Overexpression of *IPMS1*ΔR variants partially rescue the phenotype of *IPMS*-null *E. coli* strain and *Arabidopsis* mutant

Next, we reassessed the physiological consequence of C-ter deletion of IPMS in *E. coli* and *Arabidopsis*. To this, we initially performed the complementation assay in the Leu-deficient *E. coli* strain CV512 (DE3). Briefly, the pET28a constructs expressing the full-length Bju.IPMS1-A1 and Bju.IPMS1-A2 proteins, and their corresponding ΔR variants viz., Bju.IPMS1-A1ΔR and Bju.IPMS1-A2ΔR, as well as the CRISPR-mimics a_1_a_1_ΔR and a_2_a_2_ΔR (mimicking the C-ter edited Bju.IPMS1ΔR proteins), were each transformed into CV512 (DE3) and the growth pattern of the transformed cells was analyzed upon IPTG induction. The OD measurement after every 12 hrs, showed that the cells harboring the Bju.IPMS1 C-ter truncated (ΔR) variants showed a delay in reaching their steady-state growth phase in comparison to their corresponding full-length *Bju.IPMS1-A1* and -*A2,* which reached the stationary phase after 36 hours. A similar delay in the growth of ΔR variants was also observed in solid medium (**Figure 4A & 4B**).

**Figure 4.**
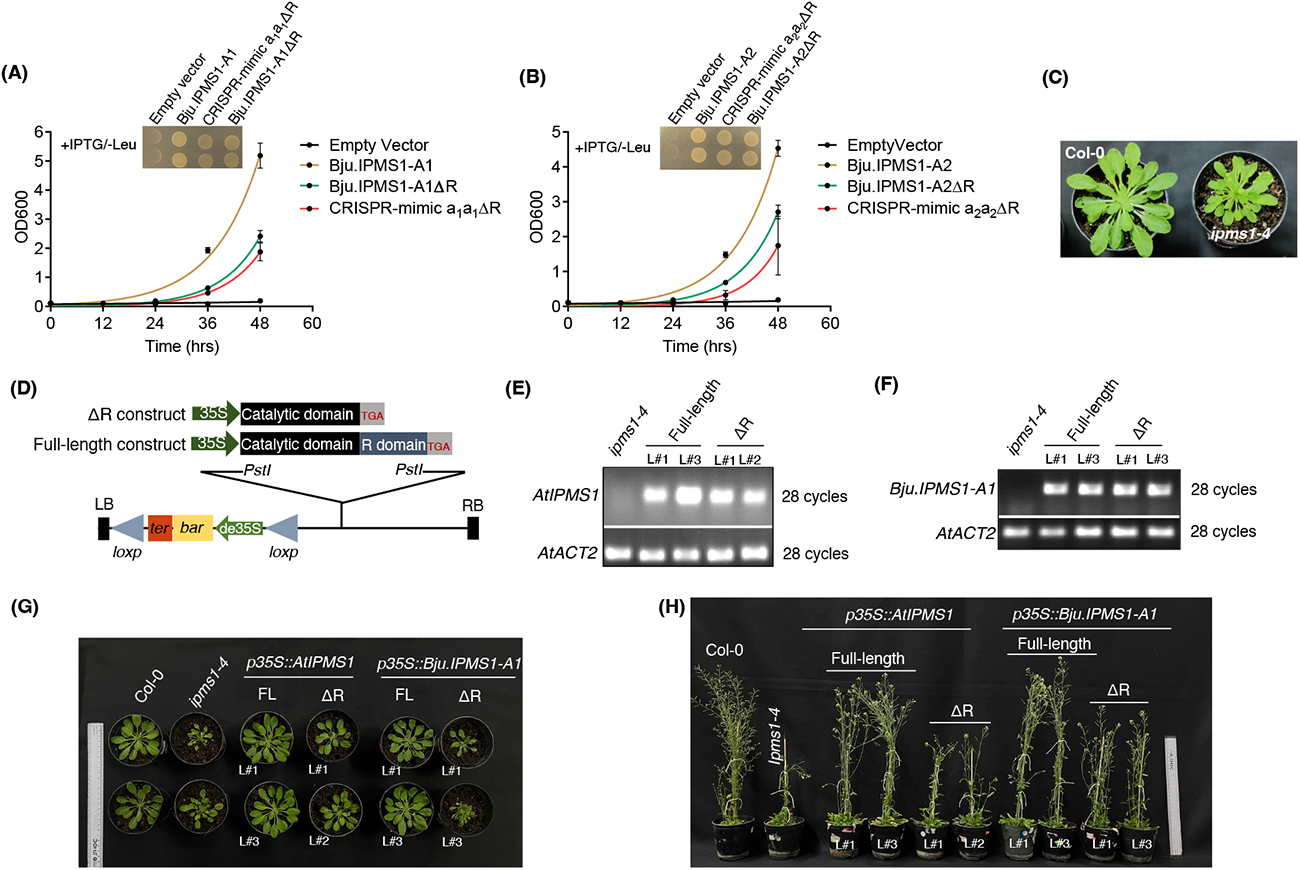
Functional complementation of *IPMS-*null *E. coli* strain CV512(DE3) and *Arabidopsis ipms1-4* mutant using C-terminal truncated variants of *Bju.IPMS1*. Growth curve analysis of CV512(DE3) complemented with pET28a constructs expressing **(A)** Bju.IPMS1-A1 and its ΔR variants; and **(B)** Bju.IPMS1-A2 and its ΔR variants, was performed in the M9 minimal media and measured for up to 48 hours. Values shown are from two independent cultures. The empty vector (pET28a) was used as the negative control. The corresponding growth in M9 minimal dropout (-Leu) agar added with 1 mM IPTG is shown in the inset (48 hours). **(C)** Rosette phenotype difference between wild-type Col-0 and *AtIPMS1* loss-of-function mutant, *ipms1-4*. **(D)** Over-expression constructs harbouring full-length *AtIPMS1, Bju.IPMS1-A1* and the truncated *AtIPMS1ΔR* and *Bju.IPMS1-A1ΔR* genes used for complementation of *ipms1-4* mutant. Transcript analysis of (E) *AtIPMS1* in *35S:AtIPMS1* and *35S:AtIPMS1ΔR* complemented *ipms1-4* lines; and (F) *Bju.IPMS1* in *35S:Bju.IPMS1-A1* and *35S:Bju.IPMS1-A1ΔR* complemented *ipms1-4* lines. Semi-quantitative RT-PCR analysis was performed for 28 cycles and the *AtACTIN2* was used as the reference control. Phenotypic analysis include **(G)** rosette growth at the one month stage and **(H)** plant height at maturity of the *ipms1-4* mutant complemented with the full-length *IPMS1* and ΔR variants (*p35S:AtIPMS1ΔR* or *p35S:Bju.IPMS1-A1ΔR*) along with Col-0 and *ipms1-4*. Photography of two independent lines is shown.

Subsequently, we complemented a recessive *Arabidopsis* T-DNA insertional mutant *ipms1-4* (**Figure 4C**) with constructs over-expressing full-length *AtIPMS1* and *Bju.IPMS1,* and their ΔR variants (**Figure 4D**). The *ipms1-4* line displayed a retarded growth and development showing reduced rosette leaves and dwarf plant height upon bolting compared to Col-0 (**Figure 4C**). The introduction of *p35S::AtIPMS1* or *p35S::Bju.IPMS1-A1* in the *ipms1-4* mutant background completely restored the phenotype of *ipms1-4* to wild-type Col-0 (**Figure 4E to H**), suggesting that *Bju.IPMS1-A1* has a conserved function as *AtIPMS1.* In contrast, the lines generated with *p35S::AtIPMS1ΔR* and *p35S::Bju.IPMS1-A1ΔR* in the *ipms1-4* background, showed only a partial rescue with a reduced rosette and shorter plant height, despite their constitutive over-expression (**Figure 4E to H**). The data suggest that while the C-ter deleted IPMS1 is functionally active, it is unable to completely rescue the IPMS1 function under both *in vivo* and *in planta* conditions.

### The C-ter edited *a_1_a_1_ΔR* and *a_2_a_2_ΔR* mutants have low Leu contents, but higher levels of the Leu pathway intermediates

While the C-ter truncated Bju.IPMS1ΔR variant displayed a higher *in vitro* 2-OIV activity compared to its full-length wild-type enzyme (**Figure 2A**), a significantly lower accumulation of the end-product Leu in *a_1_a_1_ΔR* and *a_2_a_2_ΔR* mutants prompted us to quantify the Leu pathway intermediates. We quantified all three Leu pathway intermediates viz, 2-isopropylmalate (2IPM), 3-isopropylmalate (3-IPM) and the Leu precursor, α-ketoisocaproate (KIC) formation in the leaf extracts of *a_1_a_1_ΔR* and *a_2_a_2_ΔR* mutants using LC-MS/MS and compared them with the amounts in wild-type *B. juncea* (**Figure 5; Supplementary Table S6**). To our surprise, in the absence of the feedback regulation by Leu in *a_1_a_1_ΔR* and *a_2_a_2_ΔR* mutants, a significantly higher level of Leu pathway intermediates was observed - with up to a 35% increase in 2-IPM (12.77 ng mg^-1^ and 12.53 ng mg^-1^ respectively) and 3-IPM (18.83 ng mg^-1^ and 17.13 ng mg^-1^), and up to 10% increase in KIC compared to that present in the wild-type plant (**Figure 5A to C**).

**Figure 5.**
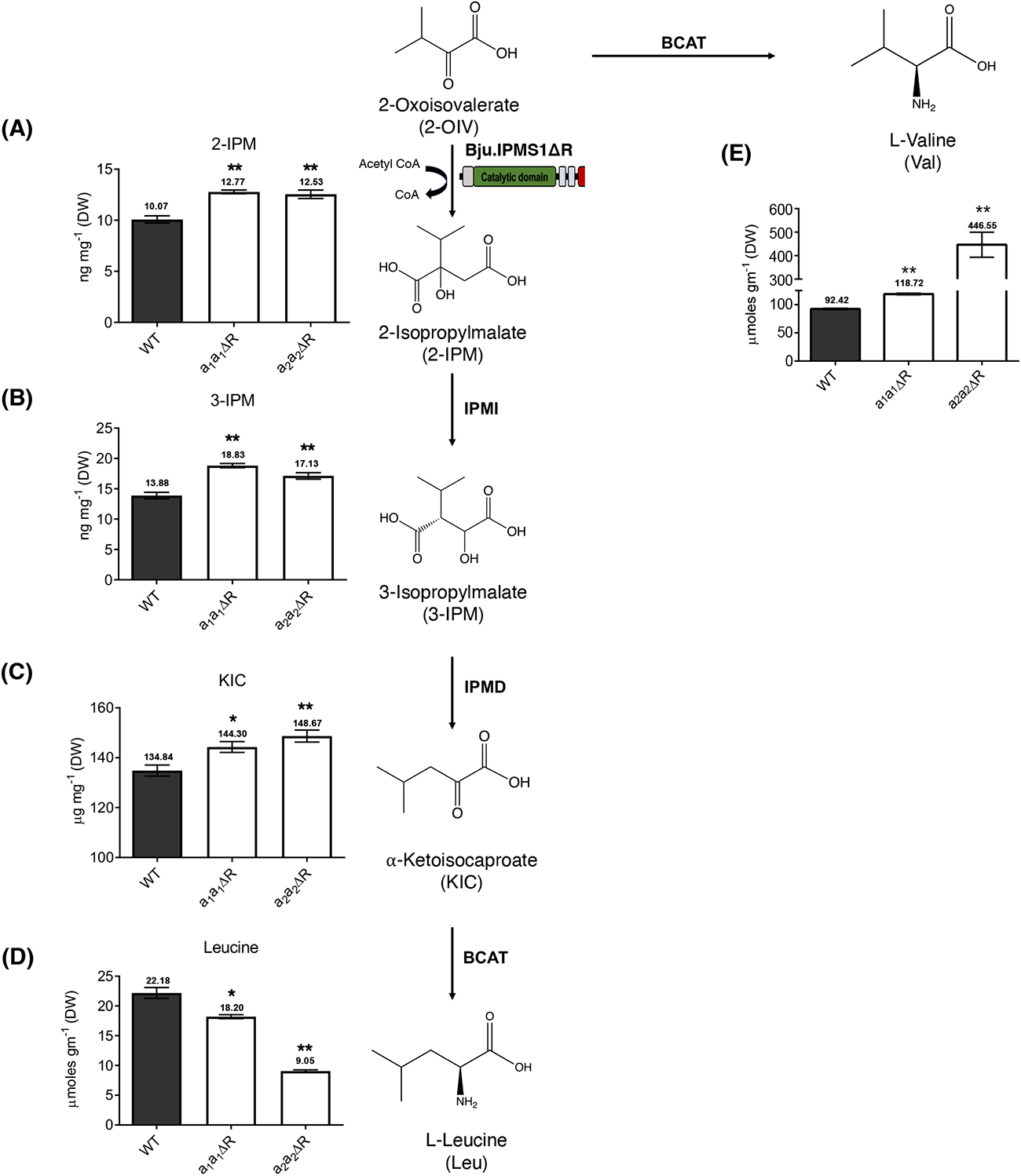
Steady-state amount of Leu pathway intermediates in the C-ter Ed Bju.IPMS1ΔR mutants. Quantification of Leu pathway intermediates in one month old leaves of *a1a1ΔR* and *a2a2ΔR* mutants along with wild-type *B. juncea* leaves using LC-MS/MS in negative ion mode (n = 6). Amounts of (**A**) 2-isopropylmalate (2-IPM), (**B**) 3-isopropylmalate (3-IPM), and (**C**) α-ketoisocaproate (KIC) in wild-type (WT) and C-ter edited mutants (*a1a1ΔR* and *a2a2ΔR*) from the T3 generation are shown. Quantifications of the Leu pathway intermediates were achieved using calibration curves developed from the pure compounds (**Supplementary Figure S5**). The reduced Leu **(D)** and increased Val **(E)** levels in the C-ter Ed Bju.IPMS1ΔR mutants are also shown in figure 3F.

The concomitant reduction in the Leu and increase in the Val levels, but an overall higher accumulation of the Leu pathway intermediates, promoted us to initially investigate the alteration in the expression of genes encoding the branched chain aminotransferase (BCAT) enzymes existing at the anabolic and catabolic pathways of the Leu and Val metabolism (**Supplementary Figure S6**). RT-qPCR analysis revealed that the transcript levels of the anabolic *Bju.BCATs* (*BCAT2*, *BCAT3*, *BCAT5*) showed no significant alteration in both *a_1_a_1_ΔR* and *a_2_a_2_ΔR* mutants in comparison to the wild-type control, suggesting that the altered Leu-Val flux is independent of the BCAT activities. Notably, the catabolic *Bju.BCAT1* gene showed a reduced transcript accumulation, specifically in the *a_2_a_2_ΔR* mutant, which could be a probable cellular mechanism to restrict the catabolism of the highly compromised Leu pool in these mutants. Hence, higher levels of Leu pathway intermediates, but decreased levels of the end-product Leu (**Figure 5D**), points to the possibility of the diversion of the Leu precursor, which in turn limits its conversion to Leu in the C-ter edited Bju.IPMS1ΔR mutants.

### Leu precursor KIC competitively inhibits Bju.IPMS1ΔR by binding to its catalytic domain

The reduced conversion rate of KIC into the end-product Leu in the *a_1_a_1_ΔR* and *a_2_a_2_ΔR* mutants led us to hypothesize that, in the absence of C-terminal regulatory domain, the catalytic domain of IPMS1 could accept KIC (4-methyl-2-oxopentanoate), which shows structural similarities with the native IPMS1 substrate, 2-OIV (3-methyl-2-oxobutanoate), but is elongated by one methylene group. To explore this possibility at the structural level, molecular docking studies on full-length Bju.IPMS1 and ΔR variants were performed with the 2-oxo acids 2-OIV and KIC using the available crystal structure of LeuA, comprising a (⍺/β)_8_ TIM barrel catalytic domain, a helical linker domain, and a regulatory domain (Koon et al., 2004). Curiously, KIC displayed a higher affinity towards the catalytic domain of the Bju.IPMS1-A1ΔR (Binding energy -4.72 Kcal/mol) compared to the full-length Bju.IPMS1-A1 (Binding energy -3.50 Kcal/mol) (**Figure 6A & B**; **Supplementary Table S7**). Both the tested proteins, however, exhibited similar binding energies with the native substrate, 2-OIV (**Supplementary Figure S7**).

**Figure 6.**
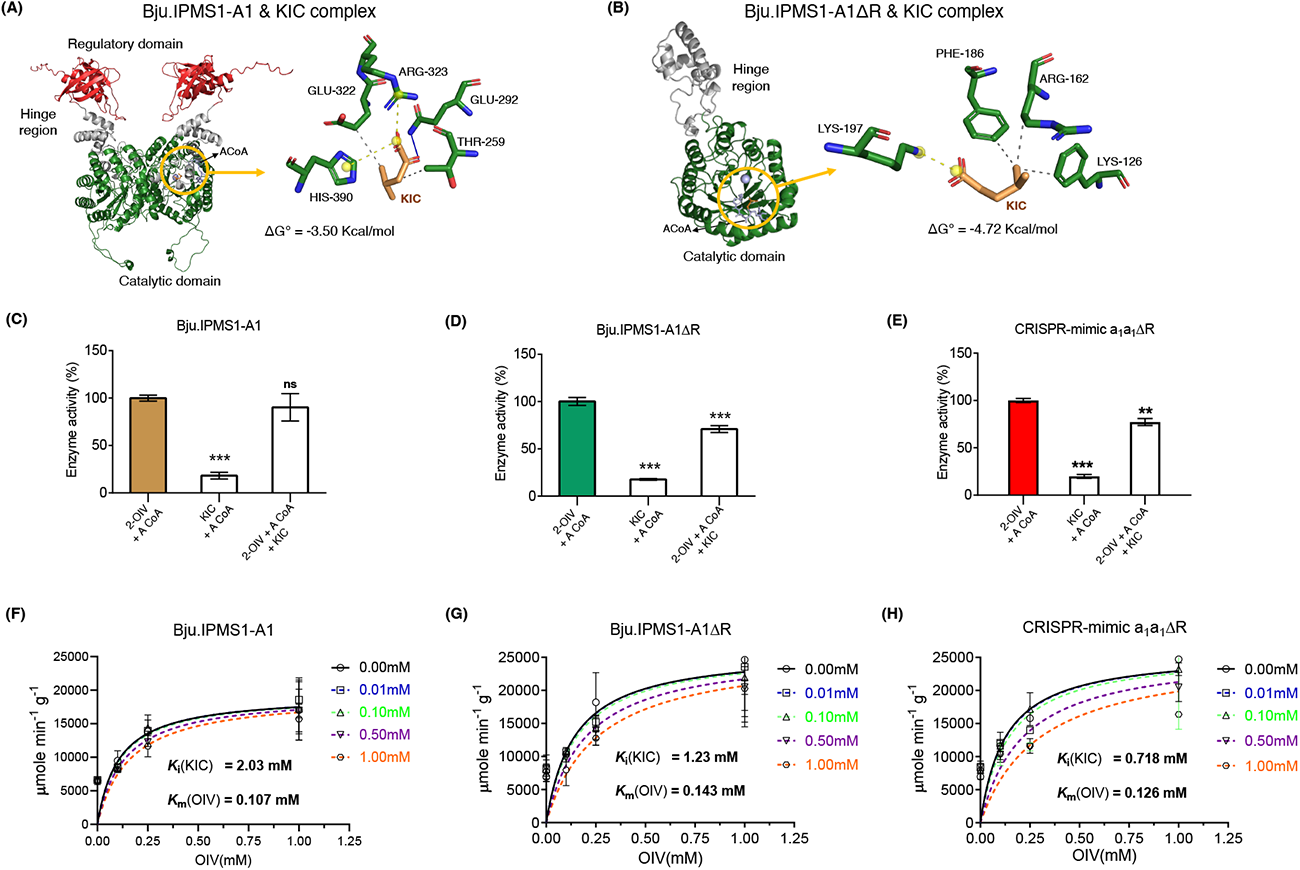
The leucine precursor 4-methyl-2-oxopentanoate (α-ketoisocaproate, KIC) competitively inhibits the activity of Bju.IPMS1ΔR variants differently. Binding position and interaction profile between KIC and **(A)** Bju.IPMS1-A1 (dimeric form) and (**B)** Bju.IPMS1-A1ΔR (monomer form). The PLIP server was used to create a two-dimensional model of the KIC-IPMS complex’s top binding position. The catalytic domain, hinge region, and regulatory domain are each depicted by a different hue in Pymol’s overall 3D cartoon depiction of the complex: Green, Grey, and Red, respectively. DTNB end-point spectrophotometric assay showing activity of **(C)** full-length Bju.IPMS1-A1, (**D**) Bju.IPMS1-A1ΔR, and (**E**) CRISPR mimic a1a1ΔR in three reaction combinations viz., (i) 2-Oxoisovalerate (2-OIV) + Acetyl CoA (ACoA); (ii) Ketoisocaproate (KIC) + ACoA; and (iii) 2-OIV + KIC + ACoA. Equimolar concentration (1 mM) of 2-OIV and KIC was used in the assay. Activities are expressed as a percentage of the activity of the respective protein in the absence of KIC. Data represents means ±SD (n=6). Kinetic assay showing the dose-dependent inhibition of (**F**) full-length Bju.IPMS1-A1, (**G**) Bju.IPMS1-A1ΔR, and (**H**) CRISPR-mimic a1a1ΔR proteins by addition of varied KIC concentrations (0 – 1.0 mM) in the presence of native substrate 2-OIV (0, 0.1, 0.25 and 1 mM) and ACoA (500 µM). Data represents means ±SD of two independent experiments.

To validate these predictions biochemically, the recombinant Bju.IPMS1-A1, Bju.IPMS1-A1ΔR and the CRISPR-mimic a_1_a_1_ΔR proteins were purified and their catalytic activities were assayed with both KIC and 2-OIV. The DTNB end-point assay was performed with (1) 2-OIV + Acetyl CoA (ACoA), (2) KIC + ACoA, and (3) 2-OIV + ACoA + KIC with 1 mM concentrations of the two 2-oxo acids (**Figure 6C to E**). The weak condensation of KIC with ACoA suggests that KIC is not a preferred substrate for either full-length Bju.IPMS1 (**Figure 6C**) or the ΔR variants (**Figure 6D & E)**. Further, when KIC was incubated along with 2-OIV, the activity of the ΔR variants was significantly reduced in comparison to 2-OIV incubation (**Figure 6D & E)**. However, the activity of WT-Bju.IPMS1 was only marginally reduced in the presence of both KIC and 2-OIV compared to that of the ΔR variants (**Figure 6C**). Further, a 2-OIV concentration at half of the maximal enzyme velocity (∼ *K*_M_), the addition of KIC in varying concentrations inhibited the activity of ΔR variants in comparison to the full-length Bju.IPMS1-A1, suggesting competitive inhibition by KIC on 2-OIV binding (**Supplementary Figure S8)**.

To get a more detailed view of this competitive inhibition, we performed a detailed kinetic assay varying five concentrations (0 to 1.0 mM) of KIC against fixed 2-OIV concentrations (0.10, 0.25 and 1 mM) in an acetyl CoA dependent manner (**Figure 6F-H**). The kinetic analysis showed that the dissociation constant (*K*_i_) for KIC towards the ΔR variants viz., Bju.IPMS1-A1ΔR and CRISPR-mimic a_1_a_1_ΔR proteins was 1.23 mM and 0.718 mM, respectively, lower than that observed for the wild-type Bju.IPMS1 enzyme (2.03 mM) (**Figure 6F**). A lower *K*_i_ value of KIC confirms its favorable binding over 2-OIV in the active pocket of Bju.IPMS1 without its regulatory domain. Hence, the absence of the C-ter regulatory domain increased the affinity of IPMS1 towards the structurally similar, but elongated inhibitor KIC compared to the native IPMS substrate 2-OIV, which in turn gives rise to a more tightly regulated Leu-independent feedback inhibition of IPMS1 activity.

## Discussion

In a metabolic pathway, feedback regulation of a rate-determining enzyme is a conserved mechanism evolved by the cellular metabolism to efficiently utilize energy and other resources (Moghe and Last., 2016; Galili et al., 2016; Farré., 2014). A well-documented example of feedback regulation occurs in the Leu biosynthesis pathway wherein the first committed enzyme, isopropylmalate synthase (IPMS), is subjected to negative regulation by Leu at its C-terminal regulatory domain. However, the significance and mechanistic basis of the IPMS C-ter regulatory domain in maintaining Leu pathway flux in plants is yet unexplored.

Here we show that the targeted editing of the IPMS C-ter regulatory domain attenuates Leu formation due to competitive feedback inhibition by the ultimate intermediate, α-ketoisocaproate (KIC) precursor. The work was performed in the crop *B. juncea,* which has a polyploid genome (AABB) and is an outcome of a natural hybridization event between diploid *B. rapa* (AA) and *B. nigra* (BB) (Augustine et al., 2014). *Brassica juncea* offers the advantage of multiple *IPMS1* homologs to manipulate the regulatory nodes involved in Leu metabolism, otherwise not available in diploid genomes. Our initial analysis suggested that each of the two progenitor genomes has contributed two duplicated *IPMS* homologs (*A1*, *A2* and *B1* and *B2*), and the four encoded *B. juncea* IPMS1 proteins (Bju.IPMS1) clustered into two distinct subclades while sharing a close evolutionary relationship with AtIPMS1 (**Figure 1B**). Furthermore, the encoded Bju.IPMS1 proteins also share conserved domain architecture containing an N-terminal (*⍺*/*β*)_8_ TIM barrel structure as the catalytic domain and a C-terminal regulatory domain similar to that reported for AtIPMS1 and *M. tuberculosis* LeuA proteins (de Kraker and Last 2017; Koon et al., 2004).

The four *Bju.IPMS1* homologs encode true functional IPMS1 enzymes with somewhat distinct biochemical parameters in the extant polyploid *B. juncea*. This was evident from the *E. coli* complementation assay wherein all four *Bju.IPMS1* proteins were able to rescue the *IPMS*-null *E. coli* mutant (**Figure 1E**), similar to that of the *Arabidopsis* and *Brassica atlantica* IPMS (de Kraker 2004; Field et al., 2006). Kinetic assays, however, revealed that the enzyme affinity (*K*_M_) of the A genome-encoding Bju.IPMS1-A1 and Bju.IPMS1-A2 proteins towards the 2-OIV substrate are higher with *K*_M_ values of 157.7 μM and 143.9 μM, respectively, than those of the B genome-encoding Bju.IPMS1-B1 and Bju.IPMS1-B2 proteins with *K*_M_ values of 216.6 μM and 230.7 μM, respectively (**Table 1**). The affinity of the Bju.IPMS1-A1 and -A2 proteins to 2-OIV were also higher than those of other plant IPMSs characterized to date (de Kraker et al., 2004; Sugimoto et al., 2021; Wang et al., 2022;), indicating that these two Bju.IPMS1 enzymes efficiently catalyse the first committed step of Leu biosynthesis in *B. juncea*.

Several plant IPMS proteins, *A. thaliana* AtIPMS1&2, *Oryza sativa* OsIPMS1&2 and *Solanum lycopersicon* SlIPMS1 have been previously demonstrated to be inhibited by Leu at their at their C-terminal regulatory domains *in vitro* (de Kraker et al., 2007; He et al., 2019; Ning et al., 2015; Xing & Last, 2017). Moreover, *in vitro* removal of the C-ter domain from AtIPMS1&2 caused insensitivity towards Leu binding and inhibition without affecting catalytic activity, unlike in MtIPMS (de Kraker & Gershenzon, 2011). Similar results were also observed for the Bju.IPMS1 (**Figure 2A & B**). Both Bju.IPMS1-A1 and Bju.IPMS1-A1ΔR could catalyze the first committed step of Leu biosynthesis with an even higher enzyme activity than the full-length protein (**Figure 2A**). In accordance with the AtIPMS1/-R protein, the Bju.IPMS1-A1ΔR variant also exhibits no feedback inhibition by the end-product Leu (**Figure 2B**) (de Kraker & Gershenzon, 2011). Thus, an *in vivo* editing of *Bju.IPMS1* regulatory region in *B. juncea* might be expected to result in a Leu biosynthetic pathway devoid of the feedback regulation on Bju.IPMS1 enzymes and thus an increased flux to Leu.

However, this was not the scenario in *B. juncea,* where the removal of C-ter regulatory domain of *Bju.IPMS1-A1* and -*A2* homologs, using CRISPR/Cas9 editing, was associated with a significant drop in the Leu content to approximately 50% of the wild-type level (**Figure 3**). These results were paralleled by the ability of IPMS1ΔR variants and the CRISPR-mimic a_1_a_1_ΔR and a_2_a_2_ΔR mutants to only partially complement the *IPMS* loss-of-function mutants of *E. coli* and *Arabidopsis* compared to the full-length *Bju.IPMS1* (**Figure 4**). This partial functional complementation points to the fact that the catalytic ability of C-ter truncated IPMS1 at the cellular level is retained, in accordance with the *in vitro* assay (**Figure 2**), but *in planta* regulation of C-ter edited Bju.IPMS1ΔR is modified in a different way with retention of some degree of apparent feedback inhibition.

Retention of feedback inhibition has also been observed in earlier studies for nucleotide triphosphate (NTP) biosynthesis in *E. coli*, where the removal of a canonical feedback regulation domain did not alter the steady state levels of the pathway end-products (UTP and CTP) by diverting excess pathway intermediate flux to uracil (Reaves et al., 2013). Metabolic analysis of Leu pathway flux in the C-ter edited Bju.IPMS1ΔR mutants with a similar loss of the Leu feedback regulation domain demonstrated that flux to the three pathway intermediates (2-IPM, 3-IPM and KIC) was retained, but the rate of KIC conversion to the end-product Leu became a rate-limiting factor for Leu accumulation. This low Leu level was simultaneously associated with high Val content, possibly due to the diversion of the Bju.IPMS1 substrate 2-OIV towards Val (**Figure 5**). The increased KIC content and an increased diversion of 2-OIV to Val can be explained by an alternative regulatory mechanism acting on the Bju.IPMS1 enzyme in the absence of its regulatory domain (C-ter Ed Bju.IPMS1ΔR).

The ultimate anabolic step and the first catabolic step and in the Leu-Val pathways are catalyzed by the anabolic and catabolic BCATs, respectively (Diebold et al., 2002; Maloney et al., 2010). However, the overexpression of anabolic *BCATs* in *A. thaliana* (Lee et al., 2018) and catabolic *BCAT* mutants in wheat (Corredor-Moreno et al., 2021) had no influence on the Leu flux. The unaltered expression of anabolic *BCATs* in Bju.IPMS1ΔR mutants also confirms the fact that the altered Leu levels, in the absence of IPMS C-ter domain, is independent of the BCAT enzymes. Furthermore, the reduced catabolic *BCAT1* expression observed in the Bju.IPMS1ΔR mutant seems to restrict the reduced Leu pool towards its catabolism (**Supplementary Figure S6**).

KIC had been previously described as a competitive inhibitor of the native IPMS in the fungus *Mortierella alpina* (Sonnabend et al., 2022). Our molecular docking analysis also showed distinct binding of KIC to both the full-length Bju.IPMS1-A1 protein and the Bju.IPMS1-A1ΔR variant. In fact, in the absence of the C-terminal regulatory domain, the N-ter (⍺/β)_8_ barrel catalytic region of Bju.IPMS1ΔR seems to have a greater affinity towards KIC than the full-length Bju.IPMS1 has (**Figure 6**). Interestingly, at the chemical level, KIC (4-methyl-2-oxopentanoate) shares a close similarity with the native IPMS substrate, 2-OIV (3-methyl-2-oxobutanoate), but with an additional methylene group in the carbon chain (**Figure 1**), thereby pointing towards KIC as a possible competitive inhibitor . This was confirmed by the *in vitro* inhibition assays where including KIC in the native IPMS reaction led to a profound inhibition of activity towards OIV for both the Bju.IPMS1ΔR and CRISPR-mimic enzymes. However, no significant inhibition was observed for the full-length Bju.IPMS1 (**Figure 6**). Taken together, the levels of Leu pathway intermediates and the *in vitro* inhibitor assays support the fact that the absence of the IPMS1 C-ter regulatory domain makes the Bju.IPMS1 catalytic domain susceptible to competitive inhibition by the intermediate immediately preceding Leu, creating a new feedback loop in the Leu pathway.

Thus, in plants, Leu biosynthesis could be potentially regulated in two different ways (**Figure 7**). The well-known conventional allosteric feedback regulation by Leu on the IPMS regulatory domain contributes to the metabolic homeostasis of Leu and Val. However, in the absence of the IPMS regulatory domain, an alternative competitive feedback regulation by the Leu pathway intermediate, KIC, is possible on the IPMS catalytic domain, and this is realized in the absence of the regulatory domain. Hence, our study suggests that the C-terminal regulatory domain may have evolved to avoid the inhibition of plant IPMSs by KIC, which would otherwise regulate Leu biosynthesis in a less accurate manner than regulation by the final product itself. This information adds to our knowledge of how particular regulatory networks have been constructed to maintain cellular homeostasis. In this case, the C-terminal regulatory domain of IPMS may be essential for accurate regulation of Leu supply. Our results suggest that efforts to engineer the Leu pathway for biofortification in crop plants must always consider the possible binding of the ultimate pathway intermediate KIC to IPMS, which could restrict the formation of Leu in unintended ways.

**Figure 7.**
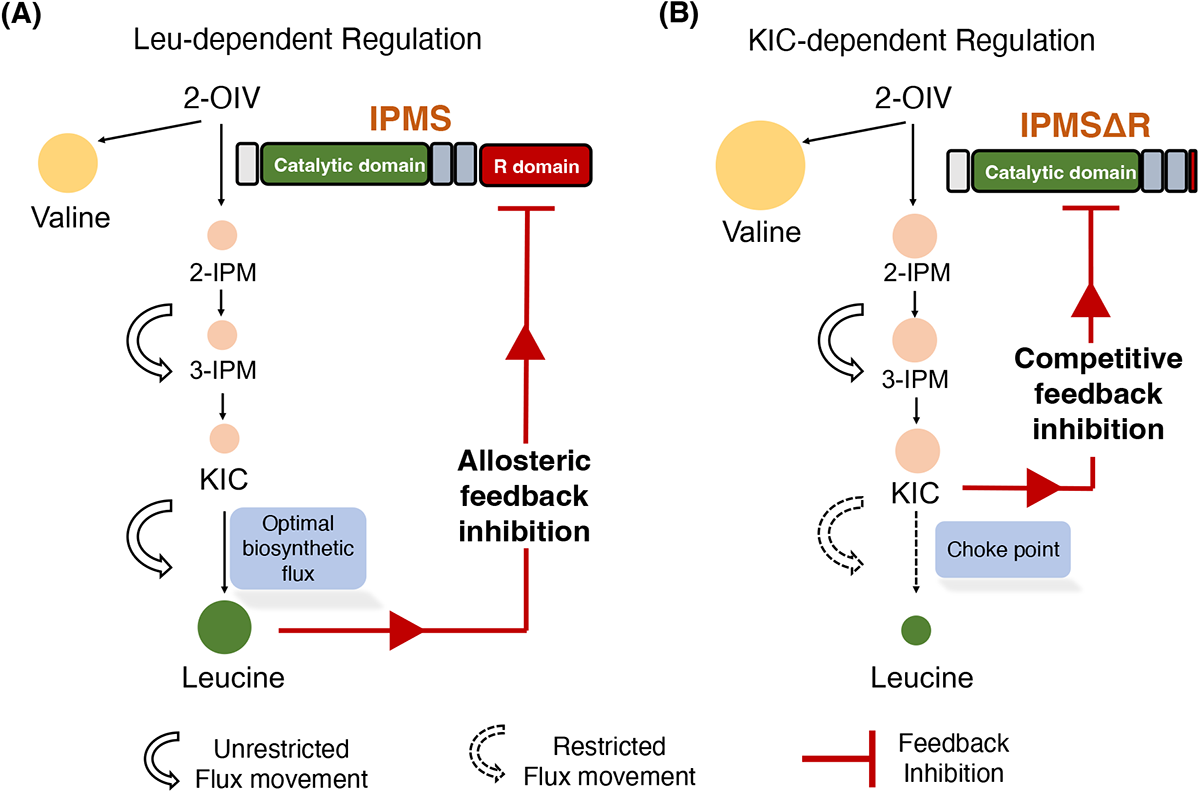
Proposed model showing the significance of IPMS C-terminal regulatory domain in maintaining the Leu homeostasis. On the left, **(A)** shows the optimal Leu homeostasis maintained by the conventional Leu-dependent feedback regulation at the IPMS regulatory domain. On the right, **(B)** shows the alternate KIC-dependent feedback regulation in the absence of the IPMS regulatory domain, at the expense of Leu homeostasis.

## Materials and Methods

### Identification and phylogenetic analysis of *Brassica IPMS1*

IPMS protein sequences of *Brassica* species were identified by performing BLASTp search using AtIPMS1 protein as a query in the BRAD v3.0 (http://brassicadb.cn) and Phytozome v13 (https://phytozome-next.jgi.doe.gov) databases using stringent cut-off values (E-value = 0). To perform the phylogenetic analysis, the identified IPMS-like protein sequences from *Brassica* species along with other functionally characterized IPMS proteins from plants, fungi and bacteria were aligned using ClustalW and the phylogenetic reconstruction was performed using the maximum likelihood method based on the JTT matrix-based model in MEGA 7 with 1000 bootstrap iterations (Kumar et al., 2016).

### Plant materials and growth conditions

Wild-type and gene-edited *B. juncea* cv. Varuna plants were grown in a controlled green-house at 24 ± 1 °C light and 18 ± 1 °C dark with 10 hr light/14 hr dark regimes and light intensity of 260-300 μmol m^−2^ s^−1^ at 60% relative humidity. Plants were grown in sand: soil rite mixture (1:2) and watered with nutrient medium twice a week. These plants were used for gene isolation, qRT-PCR analysis and for genomic DNA isolation for screening of SpCas9 induced mutations. *A. thaliana* ecotype Columbia (Col-0) and the previously described AtIPMS1 (At1g18500) T-DNA insertion mutant, *ipms1-4*, obtained from TAIR (https://www.arabidopsis.org/, Salk_101771), were grown in agropeat: vermiculite mixture (3:1) in a growth room set at 22 °C under a 16 hr light/8 hr dark cycle with 250 μmol m^-2^ s^-1^ balanced light spectrum using fluorescent lamps (40 W) at 40% relative humidity.

### Isolation of total RNA, qRT-PCR, and transcript analysis

Total RNA was isolated from different vegetative and reproductive developmental stages using Spectrum^TM^ Plant Total RNA isolation kit according to manufacturer’s instruction (Sigma-Aldrich, USA). The quality of isolated total RNA was determined in a Nanodrop (ND-1000, Thermo Scientific). Approximately, 2 μg of total RNA was reverse transcribed into cDNA with oligo(dT) primers using a First Strand cDNA Synthesis kit (Applied Biosystems, USA) in a reaction volume of 20 μL. RT-qPCR analysis of the 1:20 diluted cDNA was performed as described previously (Kumar et al., 2014). The *Actin* of *Brassica* origin was used as an internal reference control to normalize the expression data (Chandna et al. 2012) (**Supplementary Table S8**). Transcript abundance of *AtIPMS1, Bju.IPMS1* and their corresponding truncated variants in the *Arabidopsis* complemented lines were analyzed using semi-quantitative RT-PCR for 28 cycles using primers listed in Supplementary Table S8. The *AtACT2* was used as an internal reference control.

### Heterologous expression and Protein purification

For the heterologous expression of wild-type *Bju.IPMS1* proteins in *E. coli*, the CDS region, of the four *Bju.IPMS1* homologs devoid of the putative N-terminal signal sequence (Armenteros et al., 2019; TargetP 1.1 server; http://www.cbs.dtu.dk/services/TargetP/) were amplified with specific primers listed in Supplemental Table 8, and cloned within *NheI* and *NotI* sites of the pET-28a expression vector (Novagen, USA). The sequence confirmed pET-28a constructs harbouring individual wild-type *Bju.IPMS1* homologs were transformed and expressed in *E. coli* BL21(DE3) cells grown in Luria-Bertani with 50 μg ml^-1^ kanamycin. Upon an *A*_600_ of 0.6, the culture was induced with isopropyl-1-thio-β-D-galactopyranoside (1 mM final) and the cells were then grown with constant shaking at 180 rpm overnight at 18 °C. Following the centrifugation, the cells were lysed and subsequent recombinant protein purification was performed over an Ni^2+^-NTA-agarose column (4 °C) following Kumar et al. (2019).

### Generation of *Bju.IPMS1* CRISPR construct and plant transformation

The vectors for CRISPR/Cas9-mediated mutagenesis at the C-ter region of *Bju.IPMS1* homologs (*A1* and *A2*) were developed in the basic pZP200debar::SpCas9 binary vector available in the laboratory (Mann et al., 2023). The designing of 20 nt sgRNAs-1 and 2, each targeting the tenth exon of *Bju.IPMS1-A1* and -*A2* homologs and the development of AtU6-26 promoter:gRNA:scaffold expression cassette was performed as previously described (Mann et al., 2023). The gRNA1 and gRNA2 expression cassettes were cloned tandemly between *SpeI/PacI* and *AscI/MfeI,* respectively into the binary vector to develop the final pPZP200debar::SpCas9:gRNA1/2 gene editing construct. All primers used for cloning are summarized in Supplementary Table 8.

The introduction of the final gene-editing construct into the *A. tumefaciens* strain GV3101 and genetic transformation of wild-type *B. juncea* (cv Varuna) were performed as per previously described protocols from the laboratory (Augustine et al., 2013). The rooted primary transgenic T0 plants were transferred directly to soil according to the guidelines of the Department of Biotechnology, Government of India.

### Identification and molecular characterization of CRISPR/Cas9 mutant lines

Initial identification of primary transgenic plants (T0) transformed with CRISPR/Cas9 expression vector carrying *bar* resistance gene were performed by spraying with 200 mg L^-1^ Basta (glufosinate-ammonium herbicide, Bayers). T0 transformants were selfed and segregated to the advanced generations harboring stable homozygous mutations. About 50 mg of frozen leaves from T0, T1, T2 and T3 gene editing *B. juncea* plants were used for genomic DNA extraction using Cetyltrimethylammonium bromide or CTAB method (Rogers and Bendich, 1994). The extracted DNA was then used as a template to amplify the genomic region spanning the gRNA targeted site by using homolog-specific primers (**Supplementary Table S8**). To characterize the mutation at the C-ter region of *Bju.IPMS1* alleles, the gel-eluted PCR products from T0 lines were cloned into pGEM-T Easy vector (Promega, Germany) or pJET1.2 blunt end vector (Thermo Scientific™, USA) and plasmids from 10 individual colonies were Sanger-sequenced. The chromatogram from the gene-edited plants in the T2 generation, was analyzed in Synthego’s ICE (Inference of CRISPR Edits) analysis tool v3 (ice.synthego.com; Conant et al 2022). Raw Sanger sequencing reads from the *Bju.IPMS1-A1* and -*A2* C-terminal region of gene-edited mutants (.abi files) were used as query, and compared to their corresponding wild-type sequence (non-edited). The analysis showed the percentage of *Bju.IPMS1* alleles mutated at its C-ter region showing the homozygous and heterozygous nature of the CRISPR-induced mutations.

### Generation of overexpression constructs and complementation of Arabidopsis *AtIPMS1* mutant, *ipms1-4*

To generate *pPZP200:lox(bar)::35S:AtIPMS1, pPZP200:lox(bar)::35S:Bju.IPMS1-A1*, *pPZP200:lox(bar)::35S:AtIPMS1ΔR* and *pPZP200:lox(bar)::35S:Bju.IPMS1-A1ΔR* constructs, the full-length coding DNA sequences of *AtIPMS1, Bju.IPMS1-A1* and their corresponding truncated variants amplified devoid of regulatory domain were cloned initially in the entry vector pRT100 in its MCS between the *Cauliflower Mosaic Virus* 35S promoter (CaMV 35S) and poly-A tail (Topfer et al., 1987). The entire expression cassette thus created was excised and finally cloned directionally at the *PstI* double restriction sites of pPZP200:lox(bar) binary vector (Hajdukiewicz et al., 1994). Each construct was transferred into *A. tumefaciens* strain GV3101 and subsequently into the homozygous T-DNA mutant *AtIPMS1* (*ipms1-4*) genetic background by the floral dip method (Clough and Bent, 1998). The genotype of the *ipms1-4* (SALK_101771) background from ABRC was validated by PCR before floral dip transformation as described by Ajjawi et al. (2010). Homozygous T-DNA insertional mutant lines were self-pollinated and propagated. The genotyping primers are summarized in Supplementary Table 8. Complemented lines were identified using the herbicide Basta (10 mg L^-1^) with at least two independent lines for each construct and analyzed the respective transcript abundance and their phenotype.

### Site-directed mutagenesis and mutant protein analysis

To mimic the CRISPR/Cas9-induced mutations identified in the gene edited lines, site-directed mutagenesis was employed by designing primers harboring mutations similar to that observed in *a_1_a_1_ΔR* and *a_2_a_2_ΔR* mutants (**Figure 3C & D; Supplementary Table S8**). pET28a constructs expressing ORF region of the full-length *Bju.IPMS1-A1* and -*A2* were used as the template. The PCR products formed are then subjected to the *Dpn*I digestion overnight to digest the methylated parental DNA template and then transformed to *E.coli*. The fidelity of the mutation in the resulting *pET28a::CRISPR-mimic a_1_a_1_ΔR* and *pET28a::CRISPR-mimic a_2_a_2_ΔR* were confirmed by sequencing and analysis using DNASTAR software (Lasergene) and by protein purification and separation using SDS-PAGE.

### Complementation of *E. coli* CV512 (DE3)

For growth curve assay, the pET28a vector independently harboring *Bju.IPMS1-A1*, *Bju.IPMS1-A2*, *Bju.IPMS1-A1ΔR*, *Bju.IPMS1-A2ΔR*, *CRISPR-mimic a_1_a_1_ΔR* and *CRISPR-mimic a_2_a_2_ΔR* were transformed in *E. coli* leucine auxotroph, CV512 (DE3) (F^+^ leuY371; Somers et al., 1973). A fresh single colony was picked and cultured in 3 ml Luria-Bertani medium with kanamycin (50 μg mL^-1^) for 14 hr at 37 °C. 2 ml of the overnight grown cells were harvested, washed, and finally resuspended in 30 ml of M9-minimal media with OD_600_ of 0.1 containing antibiotic and 1 mM of IPTG. Subsequently, the culture was incubated at 28 °C shaker and OD_600_ was recorded after every 12 hr up to 48 hr. For plate-based growth assay, the pET28a vector independently harboring either full-length or truncated IPMS1 variants grown overnight in LB was diluted to OD_600_ of 1.0 and 10 µl of cells, suspended in M9-minimal media were dropped into the solid M9-minimal media in combinations constituting +IPTG/-Leu and -IPTG/+Leu.

### Enzyme assays

#### Discontinuous end-point enzyme assay

The spectrophotometric end-point assay using 5,5-dithiobis-(2-nitrobenzoic acid) (DTNB, Sigma Aldrich, USA) for assessing the specific activity of Bju.IPMS1 proteins was performed as previously described by de Kraker et al. (2007) with minor modifications. Briefly, 2.5 μg of the purified Bju.IPMS1 protein and its variants were incubated in a reaction mixture of 10 mM 2-oxoisovalerate, 500 mM acetyl-CoA, 4 mM MgCl_2_, and 100 mM Tris, pH 8.0 for 30 mins at 30 °C. The reaction was stopped by freezing the Eppendorf tube in liquid nitrogen for 1 min. To this, 200 μl of ethanol and 200 μl of 1 mM DTNB in 100 mM Tris pH 8.0 were added and mixed, and the reaction was incubated at room temperature for 10 mins for the free thiol group of CoA to react with DTNB forming a yellow-coloured, 3-carboxy-4-nitrothiophenol anion. The mixture was centrifuged for 15 min at 13000 rpm, and the absorbance was detected against water at 412 nm. The absorbance was corrected by subtracting the background of the identical assay mixture without the respective enzyme. To deduce the specific activity, the absorbance at OD_412_ was put in the Beer-Lamberts equation using the molar extinction coefficient of 3-carboxy-4-nitrothiophenol anion as ε_412_=14140 M^-1^ cm^-1^ (Kohlhaw, 1988; de Kraker et al., 2007).

#### Enzyme kinetics

The kinetic parameters of Bju.IPMS1 proteins towards 2-oxoisovalerate (2-OIV) were obtained by performing DTNB assay for 10 min at 30 °C in varying concentrations of 2-OIV ranging from 50 μM to 4000 μM while keeping acetyl CoA concentration fixed at 500 μM. Similarly, kinetic parameters for acetyl CoA were obtained by using varying concentrations of acetyl CoA ranging from 10 μM to 2000 μM while keeping 2-OIV concentration fixed at 1 mM. *K*_M_ and V_max_ were determined by fitting the enzyme progress curve using nonlinear regression analysis available in GraphPad Prism v6.01.

#### Leu feedback assay

DTNB assay was used to test the effect of Leu feedback on enzyme activity of the full-length IPMS and its variants following the addition of Leu at different concentrations viz., 0.25, 0.5, 0.75, 1.0, 1.5, 2.0, 2.5, and 5.0 mM. The rate of reaction on the addition of different Leu concentrations was determined as described above. Data are expressed as a percentage of the activity of the respective protein with no inhibitor (set at 100%).

#### Competitive inhibition assay

For competitive inhibition assay, DTNB assay was performed at 30 °C with fixed concentrations of 2-OIV (0.1 mM, 0.25 mM and 1 mM) while the concentration of the putative inhibitor, 4-methyl-2-oxopentanoate (KIC) ranged from 0.01 mM to 1.0 mM. The end-point and kinetic assays were performed for 30 and 10 min, respectively.

All the above-mentioned enzyme assays for each protein were performed from at least two independent protein preparations, each in duplicates.

### Quantification and analysis of amino acids and pathway intermediate contents

#### Amino acid quantification

The amino acid quantification of genome-edited *B. juncea* plants was performed following a previously described protocol (Bajpai et al., 2019). Briefly, 5 mg of freeze-dried leaves were homogenised in 500 μL of 80% methanol (Honeywell Research Chemicals, USA) and centrifuged at 13000 rpm at 4 °C for 20 min. The filter-sterilized 40 μL of supernatant was mixed with 360 μL of ^13^C, ^15^ N-labelled algal amino acid mix (Cambridge Isotope Laboratories, Inc., USA) prepared at a concentration of 10 μg of the mix per ml. This mixed solution was analysed directly in liquid chromatography coupled with a SCIEX 6500+ triple-quadruple-trap MS/MS at the NIPGR metabolomics facility. Separation of amino acid was done with a Zorbax Eclipse XDB-C_18_ column (50 × 4.6 mm, 1.8 μm, Agilent Technologies). The mobile phase comprised of solvent A (water, 0.1% formic acid) and solvent B (acetonitrile, 0.1% formic acid) with the following elution profile: 0-1 min, 3% B; 1-3.8 min, 3-50% B; 3.8-3.9 min, 50-100% B; 3.9-5 min, 100%B; 5-5.1 min, 100-3% B; and 5.1-7 min, 100% B with a flow rate of 0.7 ml min^−1^. The column temperature was maintained at 27 °C. For the detection of amino acids, the mass spectrometer was operated in positive ionization mode (MRM modus) to monitor the analyte parent ion to product ion (Bajpai et al., 2019). Analyst 1.5 software (Sciex) was used for data acquisition and processing. Individual amino acids in the sample were quantified by the respective ^13^C, ^15^N-labeled amino acid internal standard.

#### Estimation of Leu Intermediates

For the detection of 2-IPM and 3-IPM, 15 μL of methanolic extracts of the 20 mg of freeze-dried *B. juncea* leaves were directly injected into the LC-MS/MS. For the calibration curve, different concentrations of pure 2-IPM and 3-IPM (Sigma-Aldrich, USA) ranging from 0.1 ng/ml to 1000 ng/ml in high-grade water were used (**Supplementary Figure S5**). Amount of malates detected in the gene-edited mutants was calculated using the equation derived from the standard curve prepared for the respective malates.

For the detection of the Leu precursor, Ketoisocaproate (KIC), we followed the MS-probe derivatization protocol recommended by Kojima et al. (2009) with minor modifications. Briefly, 100 μL of either pure KIC (1 mg in 1 ml of pure water) or 80% methanol extract of plant tissue was reconstituted with 400 μL of water and passed through a water-equilibrated DEAE-Sephadex® A-25 column (Sigma-Aldrich, USA), followed by washing with 1 ml of water and elution of the column bound metabolites with 400 μL of 3% formic acid. Post evaporation, the metabolites were reconstituted with 75 μL of 1-propanol, 20 μL of water and 4 μL of bromocholine (MS-probe; Kojima et al., 2009) in 70% acetonitrile and 0.8 μL triethylamine. The solution was re-evaporated by incubating at 80 °C for 130 min and then kept on ice to cool down. Finally, the derivatized metabolites were reconstituted in a final volume of 50 μL of 0.05% formic acid, and 10 μL of the 1:1000 diluted derivatized sample was subjected to LC-MS/MS analysis.

Separation of malates (2-IPM and 3-IPM) and the derivatized KIC were performed using the same protocol as described above for the amino acid separation. The mass spectrometer was operated in negative ionization mode (MRM modus) to monitor the analyte parent ion to product ion (**Supplemental Table S5**). Analyst 1.5 software (Sciex) was used for data acquisition and processing.

### Homology modelling and Oligomer prediction

Homology modelling of Bju.IPMS1-A1 protein and its variants was performed using the Swiss-Model server (http://swissmodel.expasy.org/), wherein an AlphaFold v2 repository Q9C550 (chloroplastic, 2-isopropylmalate synthase 2 from *A. thaliana*) was the best template to utilize, with 87.64% sequence identity. The quality of the homologous 3D structure was assessed using the Ramachandran plot (Ramachandran Favored 94.55%). Further, the monomeric form predicted by the Swiss-model was processed using GalaxyGemini Server (Lee et al., 2013) to obtain the homo-dimeric form of the protein. Acetyl CoA and the metal ion was added to the structure.

### Molecular Docking

Using Autodock 4.2 (Morris et al., 2009), the two compounds 2-OIV and KIC were individually docked with the Bju.IPMS1-A1 dimeric and Bju.IPMS1-A1ΔR monomer forms. Docking energy was obtained by adding the Van der Waals and hydrogen bonding energies, whereas binding energy came from the Van der Waals and desolvation energies. Lamarckian Genetic Algorithm (GA) was employed for each run, consisting of 10 GA runs with a maximum of 27,000 generations, 0.02 rate of gene mutation, and 0.8 rate of crossover for each shortlisted ligand. The results were analyzed using the PLIP server (Adasme et al., 2021), and the visualization was done using PyMOL.

## Acknowledgements

The work was supported by the BT/PR25839/GET/119/102/2017 grant of Department of Biotechnology, DBT(India) to N.C.B. Financial supports from Council of Scientific & Industrial Research, CSIR (India) to M.V, NIPGR Short-term fellowship to R.K. and DBT-project grant to A.S. are acknowledged. Ms. Khusboo from NIPGR-Metabolomics facility, Central facilities of DBT-NIPGR and technical help from Mr. Amal Roul, Mr. Vinod and Mr. Raju Das during the study are greatly acknowledged.

## Author contributions

N.C.B. planned and supervised the research work. M.V. performed protein biochemistry, mutation screening, metabolite estimation, *A. thaliana* and *E. coli* complementation studies, real-time assays and data analysis. R.K. developed the gene-editing and over-expression constructs, Leu inhibition assay, contributed to analysis and discussed the data throughout the work. A.S. generated *B. juncea* transformants and helped in screening of *ipms1-4* homozygous mutants and real-time assays. A.L. performed the *in silico* modeling work. M.V. and R.K. collated the data. M.V., R.K., J.G. and N.C.B. discussed the results, and drafted the manuscript. All authors read and approved the final manuscript. NCB agrees to serve as the author responsible for contact and ensuring communication.

